# EASYSTIFF^®^, a portable and innovative device able to separately analyze each skin compartment for the evaluation of mechanical properties

**DOI:** 10.1101/2023.07.13.548841

**Authors:** Gael Runel, Jean-André Lapart, Julien Chlasta

## Abstract

Human skin is submitted to various factors leading to either extrinsic or intrinsic aging. In addition to inducing molecular and morphological changes, skin aging is characterized by modifications in skin biomechanical properties modifications. These modifications can be used as biomarkers to both evaluate the degree of advancement in skin aging but also to detect potential pathologies. Hence, it is from critical interest to be able to evaluate skin biomechanical properties to fully detect and monitor these alterations. Here we introduce a portable and innovative ^®^device, EASYSTIFF^®^, which is thought to *in vivo* measure stiffness properties of skin. Based on indentation principle, EASYSTIFF allows to identify both global stiffness of skin (all confounded cutaneous compartment) or compartmentalized stiffness, in which stiffness values of each compartment, i.e., the stratum corneum, the epidermis, the dermis and the hypodermis, are separately analyzed. Thus, EASYSTIFF is a complementary tool to 2D and 3D *in/ex vivo* model and an alternative to existing devices that measure skin mechanical properties, for assessing the effectiveness of cosmetic products, monitoring skin stiffness over time and to provide better understanding in the evolution of skin mechanical properties.

## Introduction

Human skin is a complex organ organized in a multilayer structure, particularly sensitive because in direct contact with our environment. Both intrinsic and extrinsic aging lead to hallmarks of aging, involving profound structural and functional changes^1^. These visible changes can both alter the integrity of the barrier function of skin and affect relationships between individuals^2^. Thus, it is from important public interest to develop effective cosmetic and dermatologic products that heal and reduce the signs of aging. In addition, to evaluate the efficiency of these products, adapted analytical tools need to be developed. During aging, dermis undergoes some changes including a remodeling of the extracellular matrix, that lead to the decrease in collagen and elastin content but also other specific molecules^3^. This decrease causes a size’s reduction of the papillary compartment in favor of the reticular compartment accompanied by a flattening of the papillae at the interface between the dermis and the epidermis^4^. Epidermis organization is also affected during ageing, such as modifications in the number of layers forming the stratum corneum, associated with a decrease in cell renewal of basal cells^5,6^. These biological changes affecting the skin structure, irrevocably lead to biomechanical modifications of the tissue, resulting for example, in a sagging skin or in a loss of elasticity and wrinkling^7,8^.

While biochemical and molecular changes of skin ageing have been well studied and characterized using both *in vitro* 2D/3D and *ex vivo* models, those models are not sufficient to measure and understand the biomechanical data and their evolution during aging and between individuals. Moreover, biomechanical data are more and more used as biomarkers as it has been shown that aging or pathologies can alters biomechanical properties of tissues or cells ^9–13^. Mechanical properties of the skin are the direct reflect of the state of skin over ageing or in a pathological context. Therefore, the notions of stiffness, firmness or elasticity emerge, and related data provide key and essential information about the physiological state of the whole skin or by cutaneous compartments. Indeed, we speak about skin firmness or skin elasticity to designate intrinsic characteristics due to the dermis, that can be altered in a context of hydration loss. In the same way, an alteration of the lipid content in the stratum corneum will lead to alterations of mechanical properties of skin. In all cases, measuring the stiffness of a component, whether it is a tissue, a cell or other, allows to identify these mechanical properties changes and can be translated into more general terms such as firmness or elasticity.

Thus, biomechanical *in vivo* study (i.e., on volunteers) of the different cutaneous compartments is an essential challenge to understand these mechanisms and eventually find out the appropriate cosmetic formulas. To assess that, many devices and tools have been generated with as very different principles.

In this study, we introduce a new innovative device, EASYSTIFF^®^ (BIOMECA, Lyon, France), able to extract the true stiffness data from the different skin compartments. EASYSTIFF^®^ was tested in clinical studies focused on skin aging where stiffness data from the different skin layers, stratum corneum, epidermis, dermis, and hypodermis, were extracted. Firstly, EASYSTIFF^®^ reliability was tested through a repeatability and reproducibility study (R&R). Then, results from clinical studies about intrinsic ageing and evolution of skin mechanical properties, of both forearm and crow’s feet, were reported.

## Results

### Working principle of Easystiff

EASYSTIFF^®^ is a portable measuring device responding to the need to characterize the mechanical properties of skin, related to an intrinsic or extrinsic ageing (**Fig1A**). To date, numerous devices are already available in the market which can evaluate viscoelastic properties of skin. The approaches used by these devices vary: suction, tissue deformation (by beads or air pulse). The analyzed parameter is the capacity of skin to return to its initial state after manipulation. At the opposite to these devices, EASYSTIFF^®^ monitors the resistance of the skin induced by a mechanical deformation. Indeed, its principle is based on the indentation of a probe into the sample (**Fig.1B**). Therefore, an indenter applies a pressure from the surface of the skin and monitors the force induced by the skin due to the indentation, while a position sensor records, with a 0.5μm sensitivity, the deformation provoked by the indenter. This assembly generates force-displacement curves, like a micro-indenter. Force-displacement curves are then processed using established theoretical mathematical models to obtain the apparent elastic modulus that corresponds to the stiffness of the skin. Thus, contrary to other existing measurement devices, EASYSTIFF^®^ directly measures the elastic modulus of the tissue that corresponds to the force generated by the measured tissue or compartment and thus directly analyzes the integrity of a compartment from a mechanical point of view.

**Figure 1:**
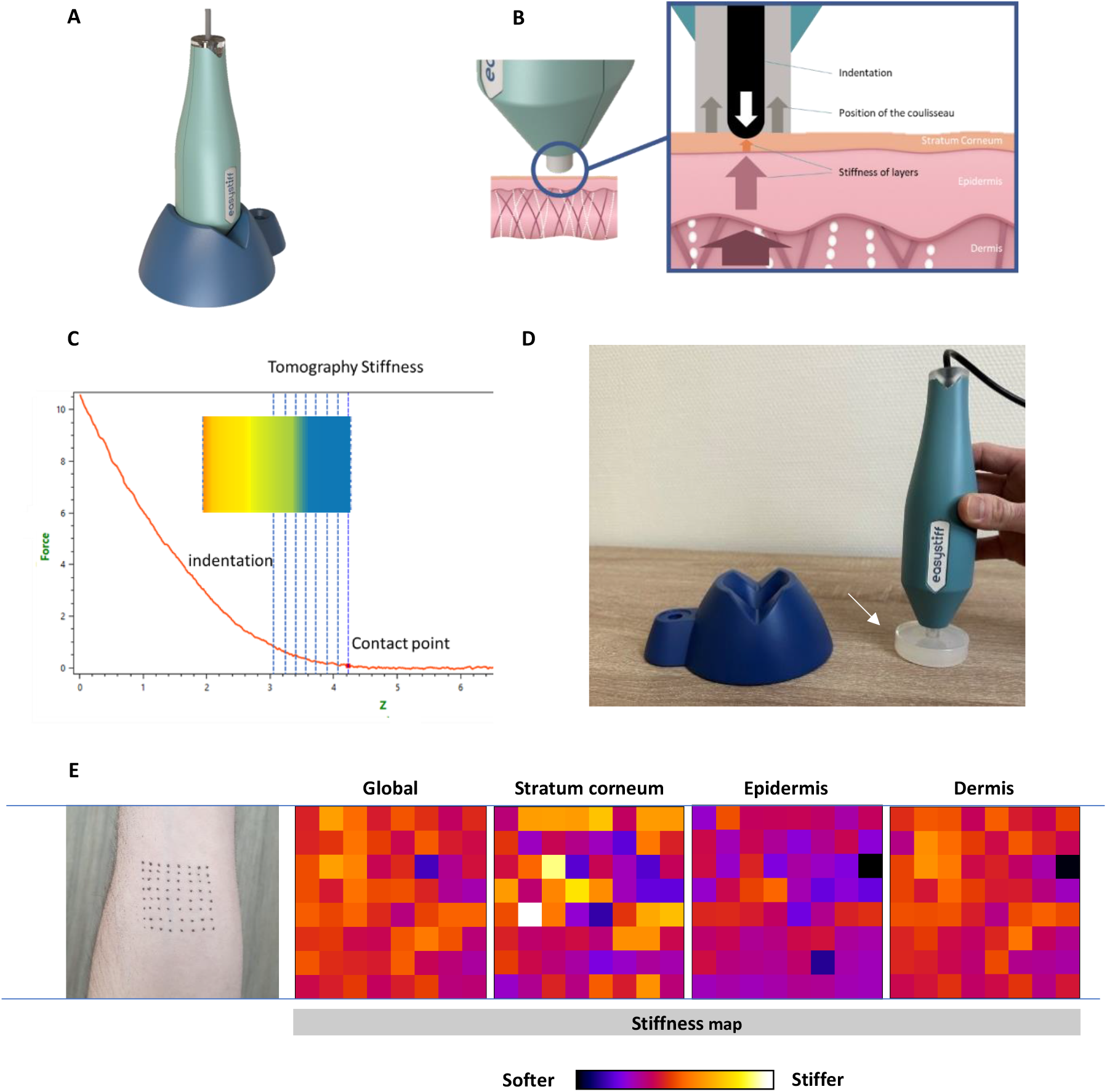
Presentation and principle of EASYSTIFF^®^. **A**) EASYSTIFF^®^ is a portable device for measuring skin stiffness. **B**) EASYSTIFF^®^’s principle is based on the generation of force-displacement curves through indentation of the probe into the sample. Concretely, a force sensor is in the probe of the device, surrounded by a slider. When measuring skin surface, the slider rises and releases the probe up to an indentation of 12mm. During acquisition, the resistance exerted by the underlying tissue is measured. **C**) The analysis of the force-displacement curves allows to generate either a global analysis to the whole skin stiffness or a compartmentalized analysis when stiffness is calculated from the data obtained by compartment. The indication of the stiffness of the sample is given by the color code (stiffer: red and softer: blue). Analysis of force-displacement curves from BioMeca analysis^®^ software. **D**) EASYSTIFF^®^ measurement test for R&R study using shore (arrow). **E**) Illustration of mechanical mapping obtained from a global and compartmentalized analysis of skin stiffness from a measurement point matrix on inner arm. The indication of the stiffness of the sample is given by the color code (stiffer: white and softer: black). Each stiffness maps correspond to the stiffness of a layer (stratum corneum, epidermis and dermis).

The main interest of this method is the possibility to monitor over time mechanical changes from a sample according to its ability to respond to different stress or cosmetic products’ effects. In addition, the extraction and analysis of force-displacement curves also allows a tomographic approach to get mechanical data, according to the compartment and depth into the tissue (**Fig.1C**). Indeed, from one hand, EASYSTIFF^®^ is able to perform a compartmentalized analysis (illustrated by vertical blue lines in **Fig1.C**) by revealing existing mechanical differences within the epidermis (ie. Stratum corneum and other layers) and in the dermis. In the other hand, EASYSTIFF^®^ allows to generate mechanical maps depending on the depth of the measured area with an axial resolution of 50 μm (**Fig.1E**). Thus, these deeper analyses allow to go further in the understanding and evaluation of the phenotypic changes of a sample.

### Repeatability and reproducibility study (R&R)

To validate the repeatability and reproducibility of the measurement by EASYSTIFF^®^, a R&R study was conducted. The aim of this approach is to measure the causes influencing the results of the measurements carried out by the EASYSTIFF^®^ device. The R&R study is a statistical method for evaluating the performance of a measurement system in terms of repeatability and reproducibility. It was performed using a protocol established in the field of ISO 17025 for the reproducibility measurement. Two EASYSTIFF^®^ devices were used and three measurements by devices were made. This was performed by four different operators, for a total of 24 measurements. Those tests aim to extract the variance of measurements according to the conditions, and to determine the reliability and the dependance of measurements in a multiparametric way. Indeed, in the R&R study, was included the analysis of the repeatability (i.e. the percentage of the total variability due to the measuring equipment), the reproducibility (i.e. the percentage of total variability due to the operators), the R&R (i.e. the percentage of total variability due to the combination of repeatability and reproducibility) and finally, the sample (i.e. the percentage of total variability due to the measured parts). The standardized ASTM D2240-15 method using specific shores was applicated in this R&R study (**Fig.1D**).

The R&R analysis allowed to determine a total variability due to measuring equipment of EASYSTIFF^®^ of 8.53%. Moreover, the percentage of total variability due to operators and due to combination of repeatability and reproducibility is respectively 2.67% and 8.94%. At last, the total variability due to measured shores is estimated to 2.57%. Given that the percentage of total variability of the different analyzed parameters is inferior to 10%, EASYSTIFF^®^ is considered as “efficient”.

### Well-Ageing clinical study about arm and crow’s feet

EASYSTIFF^®^ is a measurement device mainly dedicated to the evaluation of skin mechanical properties. It was therefore used in a clinical study where we evaluated skin stiffness from two areas of the body: the inner left arm (**Fig.2**) and the crow’s feet (**Fig.3**). This clinical study was performed with 47 volunteers between 19 to 65 years old. All volunteers declared to remove cosmetic or dermo-cosmetic product before each measurement.

**Figure 2:**
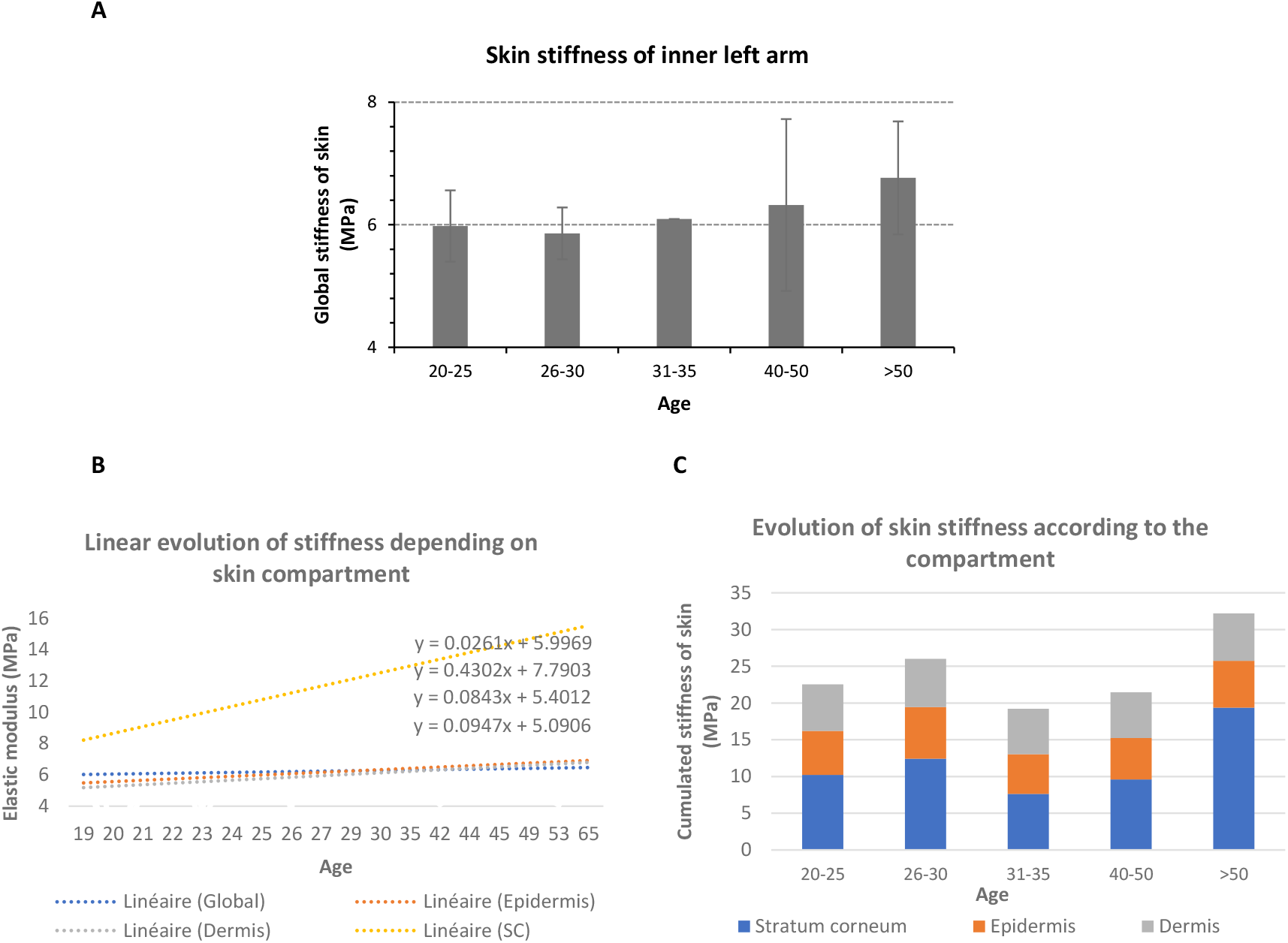
Evolution of skin stiffness from inner left arm. **A**) Skin stiffness from inner left arm classified as age range (20-25yo; 26-30yo; 31-35yo; 40-50yo; >50yo). **B**) Linear evolution of stiffness over aging of whole skin and depending on skin compartment. **C**) Evolution of skin stiffness represented as cumulative stiffness of the different compartments and classified as age range (20-25yo; 26-30yo; 31-35yo; 40-50yo; >50yo).

**Figure 3:**
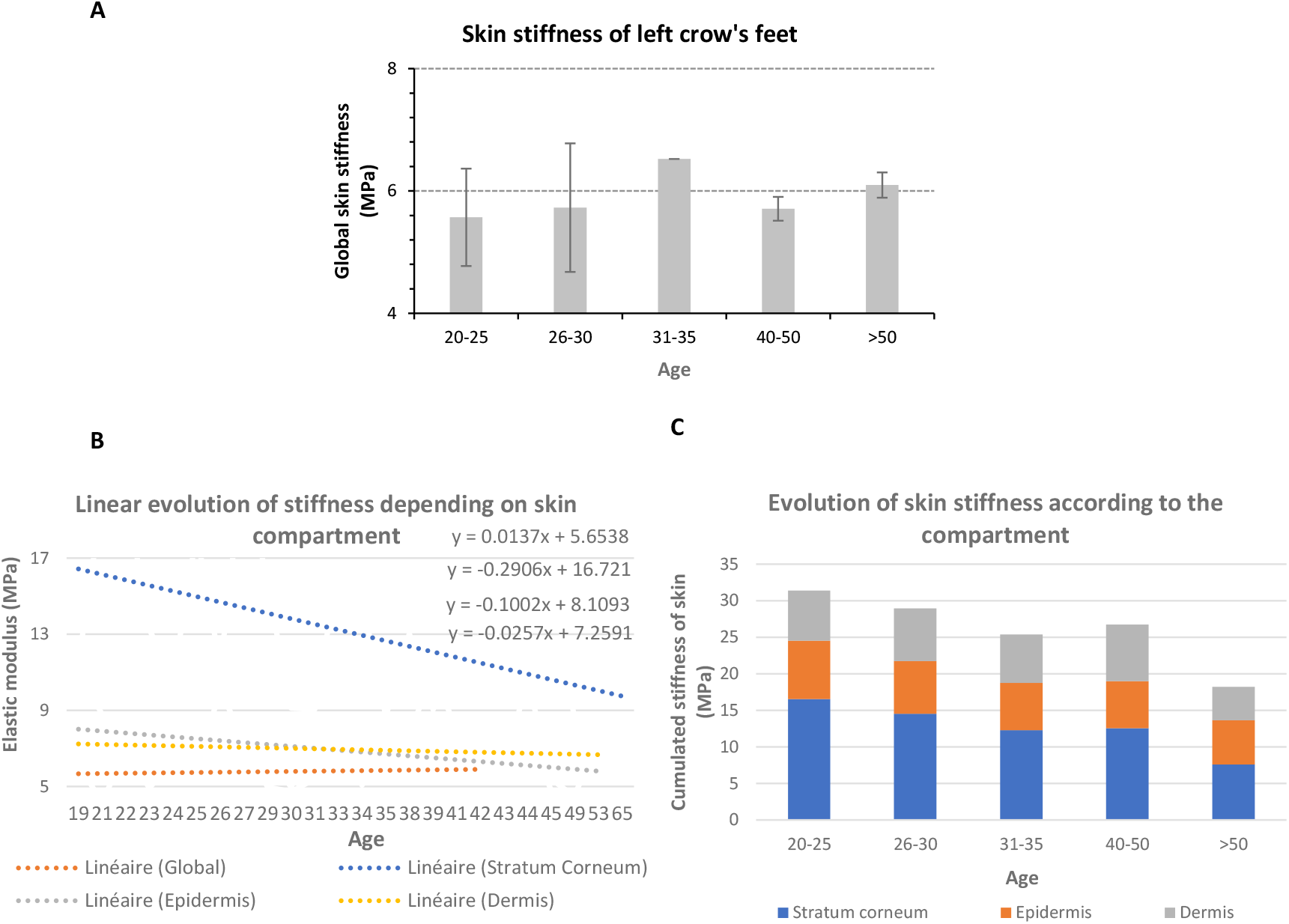
Evolution of skin stiffness from crow’s feet. **A)** Skin stiffness from crow’s feet classified as age range (20-25yo; 26-30yo; 31-35yo; 40-50yo; >50yo). **B)** Linear evolution of stiffness over aging, of whole skin and depending on skin compartment. **C**) Evolution of skin stiffness represented as cumulative stiffness of the different compartments and classified as age range (20-25yo; 26-30yo; 31-35yo; 40-50yo; >50yo).

**Figure 4:**
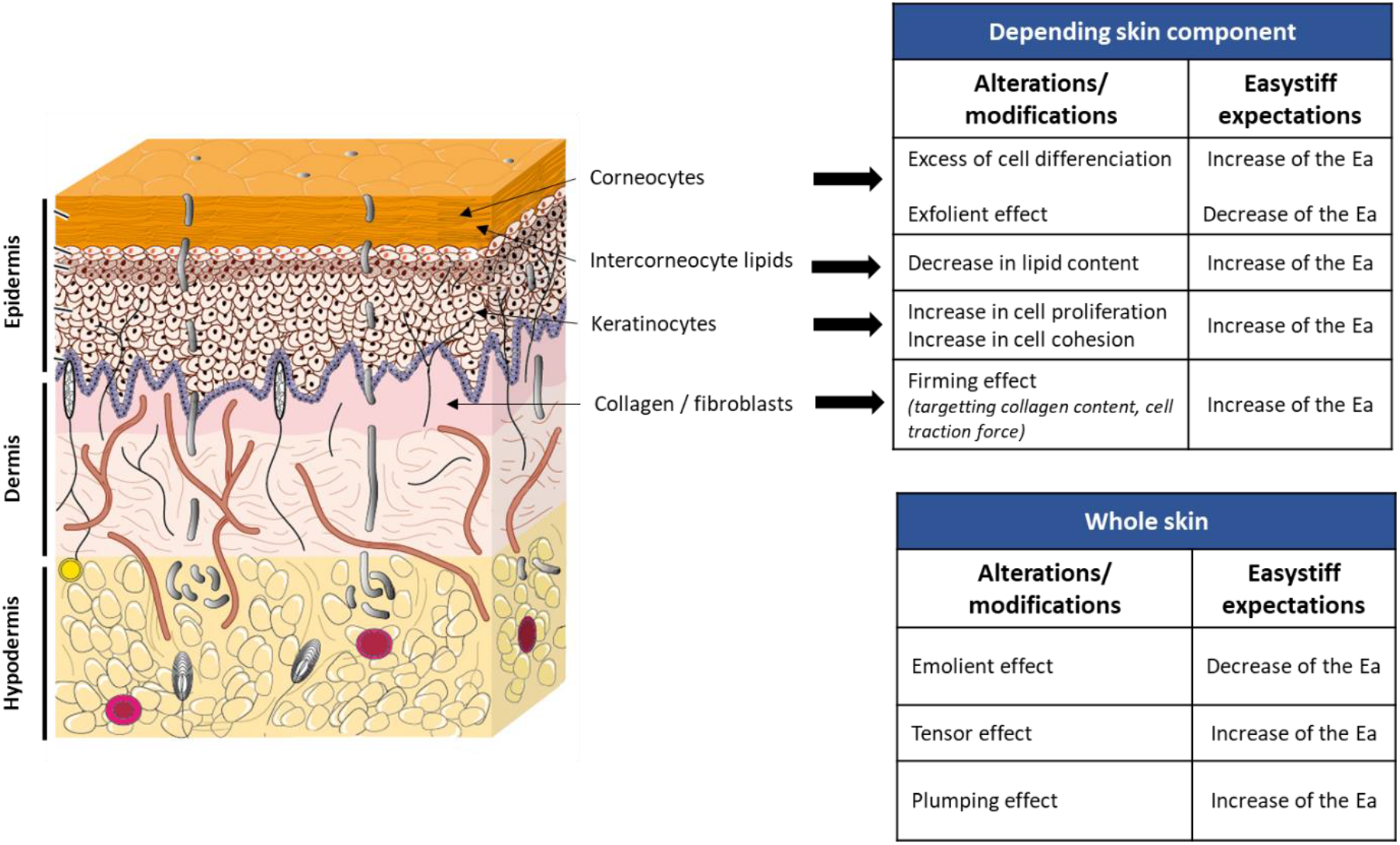
Summary of skin alterations observed by EASYSTIFF^®^.

Study of skin stiffness by EASYSTIFF^®^ revealed a progressive increase of whole skin stiffness of the inner left arm from 20-25yo to >50yo (**Fig.2A**). More precisely, the global elastic modulus of the skin appears homogeneous until 31-35yo, to then progressively increases until >50yo. Thus, whole skin stiffness from inner arm slightly increases during ageing. Moreover, an important specificity of the device is to be able to make a segmented analysis by compartment. This allows to precisely and separately analyze the stiffness of the stratum corneum, the epidermis and the dermis. Thus, we obtain more information about their stiffness evolution, that could be hidden when considering the stiffness of whole skin (**Fig.2B** and **C**). **Fig.2B** shows linear evolution of skin stiffness during ageing for each compartment.

Linear stiffness from the dermis and epidermis slightly increases in a similar pattern during aging. At the opposite, the increase of the linear stiffness of the stratum corneum is more important. This is confirmed by the **Fig.2C** where cumulated stiffness for each compartment reveals a clear increase of stratum corneum’s stiffness, compared to other compartments after 50 yo.

These data indicates that while EASYSTIFF^®^ can detect slight variations of stiffness of whole skin, it also provides important information about the variations of stiffness depending on skin compartment.

In addition to the skin of inner arm, EASYSTIFF^®^ was also used in a clinical study to evaluate the evolution of the stiffness of another part of the body, the crow’s feet, during ageing. At the opposite of the inner arm, stiffness evolution of skin from crow’s feet does not follow a progressive pattern (**Fig3.A)**. Indeed, we observed a first phase where global skin stiffness of crow’s feet tends to increase from 20-25 yo to 31-35 yo. Then a second phase is observed, where skin stiffness decreases from 31-35 to 40-50 yo to at last increasing again until >50 yo.

Moreover, the tomographic analysis performed with the device, indicated that the stiffness of the stratum corneum of crow’s feet strongly decreases during ageing (**Fig3.B)**. This is accompanied by a slight decrease of the stiffness of epidermis, while dermis stiffness is quite stable. This phenomenon is also observed in **Fig3.C** by the cumulative stiffness where a progressive decrease of the stiffness of the stratum corneum and in a less extend manner from the epidermis is observed during ageing.

Based on these results, EASYSTIFF^®^ can measure and detect variations of skin stiffness over ageing. Moreover, whole skin stiffness from crow’s feet does not evolve in a similar pattern than skin from inner arm. In the case of crow’s feet, the progressive decrease of stiffness of all compartments, more pronounced in the stratum corneum, informs on a thinning of skin and/or a sagging effect in this area. In addition, collected data show that stiffness of the stratum corneum highly varies during ageing and not necessarily in the same way, depending on the geographical area of the body.

## Discussion

In this article, we presented EASYSTIFF^®^, a new portable measuring device, as a new solution for *in vivo* assessing and analyzing the mechanical properties of skin. Skin, the first barrier of our organism protects us against aggressions and is permanently subjected to numerous alterations, whether it is due to intrinsic or extrinsic factors ^1^. These alterations lead to modifications in all skin compartments: the epidermis (including the stratum corneum) and the dermis. These modifications, firstly lead to molecular and biochemical responses from cells and tissues^14^. Then these genetic and proteomic responses are then visible at a biomechanical level. Indeed, each skin compartment possess its own range of stiffness that evolves during aging or after a specific stimulus ^8,15,16^. Thus, stiffness of cells, compartments, sub-compartments and hence tissues can be used as a skin biomarker to go further in the understanding of skin aging or to highlight skin abnormalities. These last can be benign as for example a slight dryness of skin surface or more important in the case of deeper skin damage affecting dermis and collagen network or other components of the extracellular matrix. As a result, being able to study and characterize the effect of cosmetic solutions targeting these skin disorders is necessary (**Fig 4**).

EASYSTIFF^®^ device can record stiffness of skin. Results obtained during our clinical study, reveal that EASYSTIFF^®^ can detect variations in the stiffness of skin during ageing and depending on the body area. One of the advantages of this device is that its principle is based on indentation method, in a similar principle as the one found in atomic force microscopy. Hence, this device directly records the skin stiffness, while indenting it, thanks to two sensors; a force and position sensors, included in the device. This principle is different from other existing devices that also evaluate some mechanical parameters of skin. Indeed, technologies used by these devices are based on suction or aspiration ^17,18^, ballistometry ^19^, twist ^20^ or air flow ^21^ approaches. In the case of suction, the ability of the skin to resist and its ability to return to its original position is analyzed. The principle of devices using an air flow is based on a non-contact measurement where an air flow is projected onto the surface of the skin. The cutaneous deformation and relaxation are analyzed, allowing to extract quantitative data about tension, firmness, depth, and volume of depression of the skin and the subcutaneous compartment. Device using ballistometry approach studies the impact of an object at constant and known force with skin, based on Newtonian mechanics, and measures its firmness by indentation and the dynamic resilience by the degree of rebound. At last, device using twist method relies on a mechanical twisting of the skin where the resistance and the ability of skin to return to its original position is analyzed, allowing to measure viscoelastic properties mainly for the stratum corneum but also deeper in skin. All devices using these methods have been extensively experimented for measuring the effectiveness of anti-aging products by analyzing mechanical properties of skin for dermatological and cosmetic applications. Comparative studies are leading to evaluate similarities and differences in term of accuracy, reproducibility, and efficiency, from some of these different devices with EASYSTIFF^®^. However, differences are inevitably expected due to the multiple principles of working of these tools, making difficult their comparison.

EASYSTIFF^®^, from one hand, differs from some of these existing devices by the fact that the method by indentation allows to directly evaluate whole skin stiffness from the stratum corneum to the dermis. In other hand, analysis and processing of data, allow to segment the generated raw force-indentation curves to switch from a global analysis of skin stiffness to a compartmentalized analysis of the stiffness of skin^22^. The interest is mainly to analyze separately each sub-compartment of the skin and to not introduce a bias in the analysis of whole skin stiffness due to a sub-compartment that would present some abnormalities. Thus, the usefulness is to be able to measure directly and accurately the stiffness profile of a specific sub-compartment of skin or to determine the effect of a cosmetic product targeting a specific compartment. The analyses of stiffness of skin collected from clinical studies about skin from inner arm and crow’s feet reveal that variations in skin stiffness are more detected in the stratum corneum and in a less extent manner in the epidermis and dermis, than in whole skin stiffness. This could indicate that it is more important or interesting to separately analyze the mechanical properties of each skin compartments rather than the one of whole skin, where a compensation seems to mask modifications in mechanical properties.

This instantly measurement and analysis of the stiffness of a specific compartment by EASYSTIFF^®^ can be used for monitoring overtime the effect of a cosmetic product, to validate the effect and efficiency after a topical application, but also during a determined time.

In addition to providing simple stiffness values, collected data of skin stiffness can also be processed to generate mechanical 2D mappings, that reveal and illustrate the differences in stiffness profiles of the different compartments and their evolution, depending on the depth in skin.

Finally, EASYSTIFF^®^ could also serve as a measurement device for some skin pathologies such as atopic dermatitis or psoriasis, not as a diagnosis tool but as a monitoring tool. Indeed, these kind of skin disorders are characterized by a hyperplasia resulting in the disruption of the epidermis, both at the level of the structure and morphology, and at the level of the barrier function. Consequently, the mechanical properties of epidermis are altered, and this could be detected as well as by the measurement of stiffness of the epidermis than by generating results as a stiffness mapping, where the comparison with a non-lesional skin could show different profiles. Hence, repeated measurements overtime could bring information and understanding either about the efficiency of a treatment, new outbreaks cycle of the pathology or habituation to a treatment.

To conclude, EASYSTIFF^®^ is a novel device in the dermo-cosmetic market, that directly measures, with validated repeatability and reproducibility, the mechanical properties of skin, depending on the cutaneous compartment. This device represents a real alternative, considering the tools already available, to validate and support claims for new dermo-cosmetic products. It also provides a novel modelling about cutaneous ageing from a layer-by-layer mechanical approach, associated to a better visibility about the intrinsic evolution of skin compartments in response to skin ageing. In a conceptual way, EASYSTIFF^®^ imitates and translates into real data, the sensation obtained by the finger when it meets the skin.

## Materials and Methods

### Device

The device used is EASYSTIFF^®^ (BioMeca SAS, Lyon, France) under patent number WO 2021165624 A1.

### Data acquisition

The measurements of EASYSTIFF^®^ are based on indentation method. The central indentation aperture was 5 mm in diameter with a probe size of 2mm. One force/distance cycle was performed, consisting in the displacement of the probe into the skin for 2 seconds at 1.2 mm of maximal deep, followed by the release of the device.

### Data analysis

The data processing consisted in applying a mathematical model (Hertz model) on the force/distance curves generated and recorded. Each curve was processed, and the elastic modulus was extracted according to appropriate analysis method: global, compartmentalized or by tomography analysis.

The skin layer analysis is performed using elastic modulus extractions on a determined range of distance of indentation, obtained from bibliographic data as well as data generated by BIOMECA laboratories. Validations of the deformation approach have been performed by computer simulations.

### Repeatability and reproducibility study (R&R)

Studies about the reliability of EASYSTIFF^®^ devices and operator dependency were carried out by ISO procedure on certified elastomer blocs (shore OS-3/OO (OO-300125) according to ASTM D 2240-15 standard, Hildebrand). EASYSTIFF^®^ devices used for the R&R study were BIOMECA-0-c1cy02-0010 (=devices 1) and BIOMECA-C1CY-002 (=devices 2). The conditions of tests were as follows: temperature: 20.1 °C; relative humidity: 52.0 %; number of users: 4 (=a); number of devices: 2 (=b).

Three measurements (n) per condition were performed for a total of 24 measurements (N).

The calculation of the risks was done using Fisher’s law. R&R analysis provide information about the total variability of EASYSTIFF^®^ device, variability due to operators and due to combination of repeatability and reproducibility. If the percentage calculated is inferior to 10%, the measurement process is said to be “capable”; if the percentage is found between 10 and 30%, the process is “to be improved” and if the percentage is superior to 30%, the process is said to be “not capable”.

### Ageing Study

Our study was performed with a panel of 47 volunteers between 19 to 65 years old. Two areas were measured: the inner left arm and crow’s feet. All volunteers declare to remove cosmetic or dermo-cosmetic product before measurements.

